# CysPresso: A classification model utilizing deep learning protein representations to predict recombinant expression of cysteine-dense peptides

**DOI:** 10.1101/2022.09.17.508377

**Authors:** Sébastien Ouellet, Larissa Ferguson, Angus Z. Lau, Tony K.Y. Lim

## Abstract

*Background:* Cysteine-dense peptides (CDPs) are an attractive pharmaceutical scaffold that display extreme biochemical properties, low immunogenicity, and the ability to bind targets with high affinity and selectivity. While many CDPs have potential and confirmed therapeutic uses, synthesis of CDPs is a challenge. Recent advances have made the recombinant expression of CDPs a viable alternative to chemical synthesis. Moreover, identifying CDPs that can be expressed in mammalian cells is crucial in predicting their compatibility with gene therapy and mRNA therapy. Currently, we lack the ability to identify CDPs that will express recombinantly in mammalian cells without labour intensive experimentation. To address this, we developed CysPresso, a novel machine learning model that predicts recombinant expression of CDPs based on primary sequence.

*Results:* We tested various protein representations generated by deep learning algorithms (SeqVec, proteInfer, AlphaFold2) for their suitability in predicting CDP expression and found that AlphaFold2 representations possessed the best predictive features. We then optimized the model by concatenation of AlphaFold2 representations, time series transformation with random convolutional kernels, and dataset partitioning.

*Conclusion:* Our novel model, CysPresso, is the first to successfully predict recombinant CDP expression in mammalian cells and is particularly well suited for predicting recombinant expression of knottin peptides. When preprocessing the deep learning protein representation for supervised machine learning, we found that random convolutional kernel transformation preserves more pertinent information relevant for predicting expressibility than embedding averaging. Our study showcases the applicability of deep learning-based protein representations, such as those provided by AlphaFold2, in tasks beyond structure prediction.

## Background

Cysteine-dense peptides (CDPs) are highly constrained by disulfide bonds, producing a molecular structure that confers extreme stability against proteolytic, thermal, and chemical degradation [1, 2]. As a result, there is much interest in utilizing CDPs as drug scaffolds [3]. Naturally occurring CDPs have diverse pharmacological and toxicological effects, including activities against ion channels, G-protein coupled receptors, and enzymes, as well as cytotoxic and anti-microbial properties [4–7]. One particular class of CDP, the inhibitor cysteine knot peptides—also known as knottins—has received much interest due to their confirmed and potential uses as analgesics, anthelmintics, anti-erectile dysfunction agents, neuroprotectives, antimalarials, antimicrobials, antitumour agents, protease inhibitors, toxins, insecticides, molecular imaging agents, and drug delivery vehicles [8–10].

Another reason that CDPs are of interest is their potential in drugging “undruggable” targets [11]. Identifying small molecules capable of disrupting protein-protein interactions has proven to be a challenge, leaving many protein aggregation diseases untreated. While antibodies are capable of disrupting protein-protein interactions, their large size limits their ability to penetrate tissues and access intracellular targets. CDPs, on the other hand, are small enough to avoid these limitations while still being able to interfere with protein-protein interactions [12, 13].

One of the challenges of using CDPs for therapeutic purposes is that they are difficult to synthesize and express. While the disulfide bonds are important for the stability of the molecule, incorrect linkage can lead to misfolded non-native isomers [14–16]. Recent work using high-throughput expression systems to express CDPs in mammalian cells has allowed the screening of hundreds of CDPs for expression [17, 18]. Though this work has led to the identification of a number of CDPs that can be efficiently produced through recombinant expression, our ability to discern the expressibility of CDPs based on amino acid sequences is currently limited, necessitating prior experimentation. The ability to determine which CDPs will express in mammalian cells would be valuable in identifying those compatible with biologic production, gene therapy, and mRNA therapy, and would facilitate the design and development of recombinant CDPs for therapeutic use.

To bridge this gap, we developed CysPresso, a machine learning classifier that predicts whether a CDP will be expressible based on its primary sequence. We tested various protein representations to identify the ideal model to utilize with machine learning algorithms to predict expressibility from primary sequences. We tested the performance of representations from an ELMo language model trained on protein sequences (SeqVec) [19], a deep dilated convolutional network trained on protein functions (ProteInfer) [20], and a neural network trained to predict protein structure (AlphaFold2) [21], and determined that AlphaFold2 representations performed best at predicting expressibility. The AlphaFold2 algorithm generates four protein representations, and we found that combining the representations enhanced model performance. Stratification of the dataset into knottin and non-knottin CDPs improved predictions of expressibility for knottin peptides. Additionally, model performance for knottin peptide expressibility could be further enhanced through application of a time series classification method implementing random convolutional kernels [22]. The performance of the final model, CysPresso, evaluated by receiver operating characteristic area under the curve (AUC) yielded 0.798 for non-knottin CDPs and 0.852 for knottin CDPs. To our knowledge, this is the first study to demonstrate that prediction of recombinant protein expression in mammalian cells can be carried out successfully by supervised machine learning. This method will be useful in identifying recombinant peptides and proteins that are expressible in mammalian cells, enabling the prediction of proteins compatible with biologic production, gene therapy and mRNA therapy.

## Methods

### Data preprocessing

This study utilized previously published data [17]. In this dataset, CDPs of 30 to 50 amino acids in length were tested for expression using the Daedalus lentivirus production system in HEK293 cells and subsequently purified by reverse phase chromatography (RPC). Products were deemed successfully expressed if the RPC peak demonstrated a single peak under both oxidizing and reducing conditions, which is characteristic of a single folding state and the absence of heterogeneity.

The original dataset contained a list of 1249 CDPs, including UniProt accession number, primary sequence, source organism, and expressibility. Twenty two duplicate entries were removed, leaving 1227 CDPs (Supplemental Table 1). CDPs were identified as knottins if the keyword knottin (KW-0960) was mapped to its UniProt entry on the UniProt database [23].

### Protein representations

Protein representations, or embeddings, encode proteins into a numeric format compatible with mathematical operations used in machine learning. Representations from SeqVec were generated using the pre-trained UniRef50 ELMo model [19], resulting in 1024 per-residue features derived from the final hidden layer of the neural network. Averaging within-feature yielded an embedding of 1024 features per CDP. Similarly, representations from proteInfer were generated using the Swiss-Prot EC model [24] to yield 1100 per-residue features, which were then averaged within-feature to yield an embedding of 1100 features per CDP.

To generate AlphaFold2 representations, AlphaFold2 [21] was run using the complete database setting for Multiple Sequence Alignment (MSA) search on the Compute Canada supercomputer cluster. The five models (monomer_casp14, model weights: v2.2.0 with 8x ensembling and relaxation) that were used during the 14th community wide experiment on the Critical Assessment of techniques for protein Structure Prediction (CASP14) were utilized to generate representations, but only the representation from the model with the highest average predicted Local Distance Difference Test (pLDDT) was used for downstream predictions. Embeddings from the final hidden layer of the single (384 per-residue features), pair (128 per-residue features after averaging the first dimension), MSA (256 per-residue features) and structure module (384 per-residue features) representations were collected. To create the combined AlphaFold2 representation, the single, pair, MSA and structure module features were concatenated, producing feature vectors of 1152 dimensions. When used to train random forest algorithms, per-residue features of the AlphaFold2 representations were averaged.

### Supervised learning algorithms

As random forest classifiers perform implicit feature selection and provide a measure of feature importance [25], they were implemented through scikit-learn [26] to allow comparisons between the quality of protein representations for the prediction of expressibility. Hyperparameters were left at default scikit-learn values, other than the n_estimators (number of trees) value which was set to 300. To estimate performance between different protein models, 10-fold cross validation was used, and the same shuffle was utilized across the different protein models. When data stratification was carried out, shuffling the train and test folds was balanced to contain an equal number of expressing and non-expressing non-knottins and knottins among each fold (Table 1). When time series transformation was carried out, such as when implementing the final CysPresso model, L2-regularized logistic regression was used for supervised learning as described below.

**Table 1.** Expressibility of recombinant non-knottin and knottin CDPs.

|  | Successful<br>expression | Unsuccessful<br>expression | Total |
| --- | --- | --- | --- |
| Non-knottin CDPs | 513 (58.6%) | 363 (41.4%) | 876 |
| Knottin CDPs | 165 (47.0%) | 186 (53.0%) | 351 |
|  |  |  | 1227 |

### Random Convolutional Kernel Transformation

Embeddings from deep learning protein representations are commonly averaged before machine learning tasks are carried out [19, 27, 28]. We hypothesized that employing other methodologies such as utilizing convolutional kernels may provide better feature extraction. RandOm Convolutional KErnel Transform (ROCKET) is an approach developed for time series classification tasks that uses random convolutional kernels to transform sequential features, and is useful for capturing discriminating signals and patterns in ordered data with a low computational requirement [22]. To apply ROCKET, we created a concatenation of the four AlphaFold2 representations without any per-residue averaging. To ensure uniformity in representation size, representations of CDPs with fewer than 50 amino acids were padded with zeros. As a result, each CDP was represented as 50 residues x 1152 per-residue hidden layer neural activations. Representations were then transformed by ROCKET, convolving 10,000 randomly generated kernels over the representation, producing a new representation of 20,000 features per CDP. The ROCKET representations were then classified using a L2-regularized logistic regression model, which was chosen because linear classifiers, such as logistic regression, are well-suited for utilizing limited information from a vast number of features, and regularization plays a crucial role in preventing overfitting when the number of features exceeds the number of training examples [29]. The original ROCKET methodology also promotes the use of L2-regularized logistic regression for high classification accuracy when computational expense does not need to be minimized [17].

### Comparison of average pLDDT values

AlphaFold2 offers a confidence metric on a per-residue basis, which is referred to as pLDDT [21]. The pLDDT values of AlphaFold2 for each CDP were averaged over the entire peptide and then displayed using a violin plot. Comparisons of average pLDDT values were made using the Mann Whitney test.

### SHAP analysis

To investigate the impact of AlphaFold2 representation types and residue positions on the predictions, we trained a random forest classifier on representations of size 50×1152 per CDP and analysed individual predictions over the dataset using SHapley Additive exPlanations (SHAP), a method developed to compute the direct contributions from each feature toward the final class probability [22]. The absolute values of these contributions were summed and visualized to determine which features contribute the most towards expressibility. Utilizing SHAP values, we were able to isolate the contributions per AlphaFold2 representation type as well as the contributions per residue position. To account for differences in sequence length, SHAP values were normalized by the number of sequences of that length. As data transformation (such as ROCKET) breaks direct relationships between residue positions and feature contributions, SHAP analysis was only applied to the random forest model.

### Leave-one-out cross validation

To evaluate the quality of the classification in terms of class-specific accuracies, we computed confusion matrices for both knottin and non-knottin models using ROCKET time-series classification with the combined AlphaFold2 representations as described above using a L2-regularized logistic regression model with a decision threshold of 0.5. A leave-one-out cross validation scheme was followed, training as many model instances as samples per set and aggregating the predicted labels over all samples. We also aggregated the confidence estimates of the models to calculate a general AUC over the entire set.

### Comparisons of model performance

The models discussed in this paper (except SeqVec and UniRep-RF) were evaluated with a permutation test over 50 random permutations of the stratified dataset, with a 90-10 split for training and validation sets. The different models were ranked in order of AUC, and the average rank out of 50 permutations was determined and plotted on a critical difference diagram using previously described methods [30], with the significance level set to 0.05.

To assess CysPresso in relation to state-of-the-art models of recombinant protein expression, we conducted a comparison to UniRep-RF, a predictive model of recombinant protein expression in B. subtilis [27]. Utilizing the CDPs in this dataset, UniRep protein representations were obtained [28] and UniRep-RF was implemented according to previously established methods with hyperparameter optimization [27]. UniRep-RF was compared to CysPresso by two-tailed permutation test on AUC values with significance level set to 0.05. AUC was calculated by 10-fold cross validation and iterated over 10,000 random permutations.

### Code

Code used to generate this work is available on github at https://github.com/Zebreu/cyspresso.

## Results

### Peptide expressibility database

The library used in this study characterized the expression of 1227 CDPs [17]. The lengths of the CDPs varied between 30 to 50 amino acids and were derived from a wide range of organisms, including mammals, arachnids, amphibians, reptiles, insects, birds, molluscs, fish, plants, and fungi. Of these CDPs, 351 were classified as knottins according to the UniProt database (Table 1). The data was slightly skewed, with 58.6% and 47% of the non-knottin and knottin CDPs deemed expressible in HEK293 cells, respectively.

Using a high-throughput expression system in HEK293 cells, the expressibility of 1227 CDPs was determined [17]. Entries were cross referenced with the UniProt database to distinguish knottins from non-knottins.

### Protein representation performance

Protein representations of CDPs were generated by obtaining neural embeddings from SeqVec, proteInfer, and AlphaFold2–models that are distinct in how they generate representations. For instance, SeqVec generates protein representations by utilizing the bi-directional language model ‘Embeddings from Language Models’ to identify relationships based on sequence syntax [19]. On the other hand, proteInfer [20] and AlphaFold2 [21] are neural network models trained to predict protein function and structure, respectively. Embeddings from these models were averaged and used as inputs for random forest algorithms to predict expressibility and tested using a stratified 10-fold cross validation. AUC was used as the evaluation metric (Fig. 1A). As random forest algorithms perform implicit feature selection, the performance of the random forest algorithm becomes a measure of feature performance. Thus, we examined the performance of protein representations generated from primary sequences at predicting expressibility. Out of the different protein representations tested, SeqVec had the least predictive power but still yielded usable models with AUC averaging 0.665. When classifiers were trained using embeddings from proteInfer, performance was improved with classifier AUC averaging 0.696. Interestingly, the best performance was obtained with embeddings from AlphaFold2. By using the single representation from AlphaFold2, AUC averaged 0.781 (Fig. 1B).

**Fig 1.**
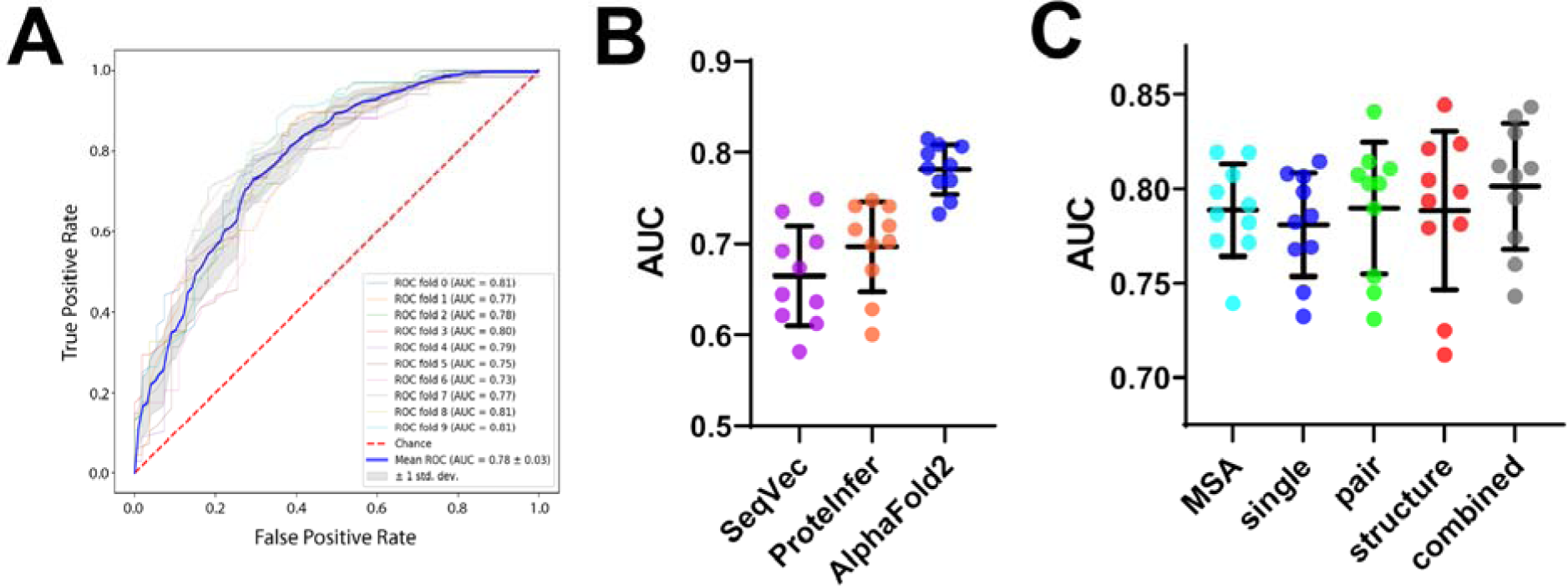
Random forest classifiers trained on SeqVec, proteInfer, and AlphaFold2 protein representations predict recombinant CDP expressibility. Random forest classifiers were trained using protein representations generated by primary sequences of CDPs and performance was estimated by 10-fold cross validation. AUC was used as the performance metric. **(A)** An example showing receiver operating characteristic curves generated using the single representation from AlphaFold2 neural embeddings. **(B)** The performance of protein representations generated by SeqVec, proteInfer, and AlphaFold2. AlphaFold2 protein representations had the highest predictive performance. **(C)** The predictive performance of neural embeddings from the four representations generated by AlphaFold2 and a concatenated combined representation. The combined representation produced classifiers with the best performance at predicting recombinant CDP expressibility. Error bars represent standard deviation of the mean value.

AlphaFold2 uses four different types of representations for structure predictions [21]. The MSA representation is based on the primary sequence and evolutionary relationships between proteins. The abstract single representation is derived from a linear projection of the first row of the MSA representation. The pair representation is a model of amino acids which are likely in close proximity based on a structure database search. These three representations are updated by the neural network and used as inputs to generate the structure module representation, which is a prediction of atom coordinates and a per-residue confidence measure. As the four AlphaFold2 representations contain different information (evolutionary relationships, abstract features, known protein structures, and atomic positions), we examined whether any of them provided superior model performance at predicting CDP expressibility, and found that AUC for MSA, single, pair, and structure module representations were similar (0.789, 0.781, 0.790, and 0.788 respectively). We then examined the effect of concatenating the four representations and observed an improvement in model performance (AUC = 0.801) (Fig. 1C).

### Generating a state-of-the-art classifier of recombinant non-knottin and knottin CDP expressibility

Having determined that combined AlphaFold2 protein representations provided the best performance for predicting recombinant CDP expressibility, we then sought to optimize the classification model. To accomplish this, we first examined whether the classifier would benefit from partitioning the data into non-knottin CDPs and knottin CDPs. Intriguingly, when the dataset was subset to knottins, the classifier performance was enhanced, although such an improvement was not observed when the dataset was subset to non-knottin CDPs (Fig. 2A).

**Fig 2.**
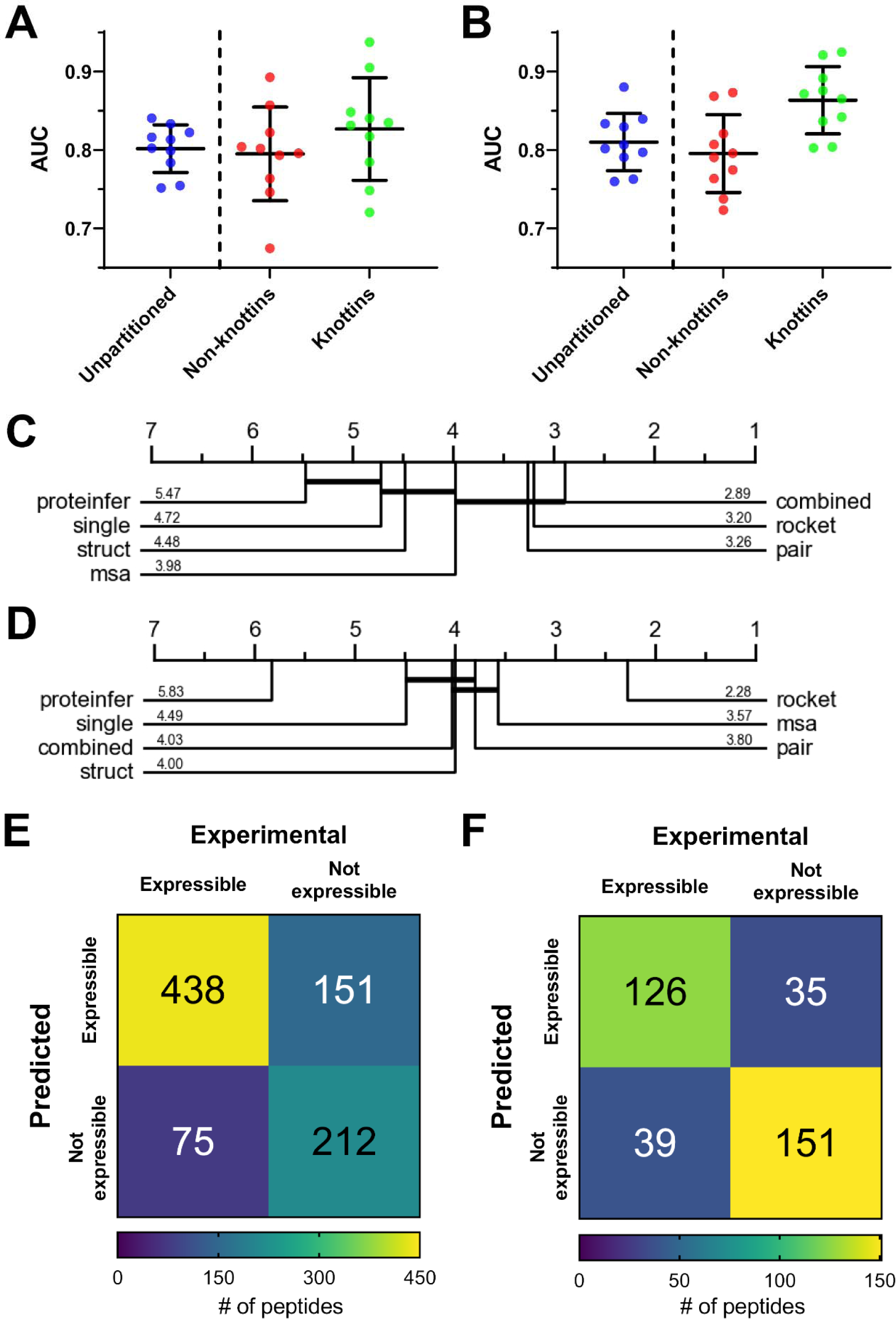
Dataset partitioning and time series classification improves prediction of knottin expressibility. **(A)** Splitting the dataset into non-knottin and knottin partitions improves prediction of knottin but not non-knottin CDP expressibility when using random forest classifiers. **(B)** Transforming AlphaFold2 protein representations with ROCKET time-series classification further improves prediction of knottin but not non-knottin CDP expressibility. Error bars represent standard deviation of the mean. **(C)** Mean AUC rank for various models at predicting non-knottin expressibility (50 permutations). Training random forest algorithms on the combined AlphaFold2 representation provided the best performance. Ranks that are not significantly different are connected by horizontal lines. **(D)** Mean AUC rank for various models at predicting knottin expressibility (50 permutations). Utilizing ROCKET on the combined AlphaFold2 representation provided the best performance. Ranks that are not significantly different are connected by horizontal lines. **(E)** Confusion matrix of the final machine learning model for non-knottin CDPs evaluated by leave-one-out cross validation. **(F)** Confusion matrix of the final machine learning model for knottins evaluated by leave-one-out cross validation.

To further improve classifier performance, a time series classification method was utilized. The neural embeddings generated by the models used in this study produce high dimensional feature vectors describing each residue in the peptide. The features were initially averaged to compress the sequence of amino acid residue features into a summary representation of a fixed size per peptide that was then used to train the random forest classifiers. While averaging preserves information from each residue, information such as order and sequential relationships are lost. Therefore, we hypothesized that by implementing a large number of random convolutional kernels, relevant features for expressibility may be captured. As a result, we applied a time series transformation technique that utilizes random convolutional kernels, named ROCKET [22], to the combined AlphaFold2 representations. Before implementing ROCKET, classifier performance (AUC) was estimated to be 0.827 ± 0.065 (Fig. 2A). Using the same cross validation folds, we found that employing ROCKET improved classifier performance for knottins and resulted in an estimated AUC of 0.865 ± 0.043 (Fig. 2B). Curiously, while the prediction of knottin expressibility was improved by the use of ROCKET, it appeared to have no impact on the prediction of non-knottin expressibility (Fig. 2B).

The proteInfer, individual AlphaFold2, combined AlphaFold2, and ROCKET models were compared by permutation test. Model performance was ranked according to AUC. For predicting the expressibility of non-knottin CDPs, the top ranked model was the combined AlphaFold2 representation without ROCKET time series transformation, though it was not ranked significantly better than the MSA, pair or ROCKET models (Figure 2C). On the other hand, for predicting the expressibility of knottin CDPs, the top ranked model was the combined AlphaFold2 representation with ROCKET time series transformation, which ranked significantly better than all of the other models (Figure 2D).

The finalized model, CysPresso, which utilizes partitioning and random convolutional kernel transformation, was evaluated by leave-one-out-cross-validation. Using this approach, classifier AUC was determined to be 0.798 for non-knottin CDPs (Figure 2E, Table 2) and 0.852 for knottins (Figure 2F, Table 2). A diagram of the machine learning model architecture is shown in Figure 3.

**Fig 3.**
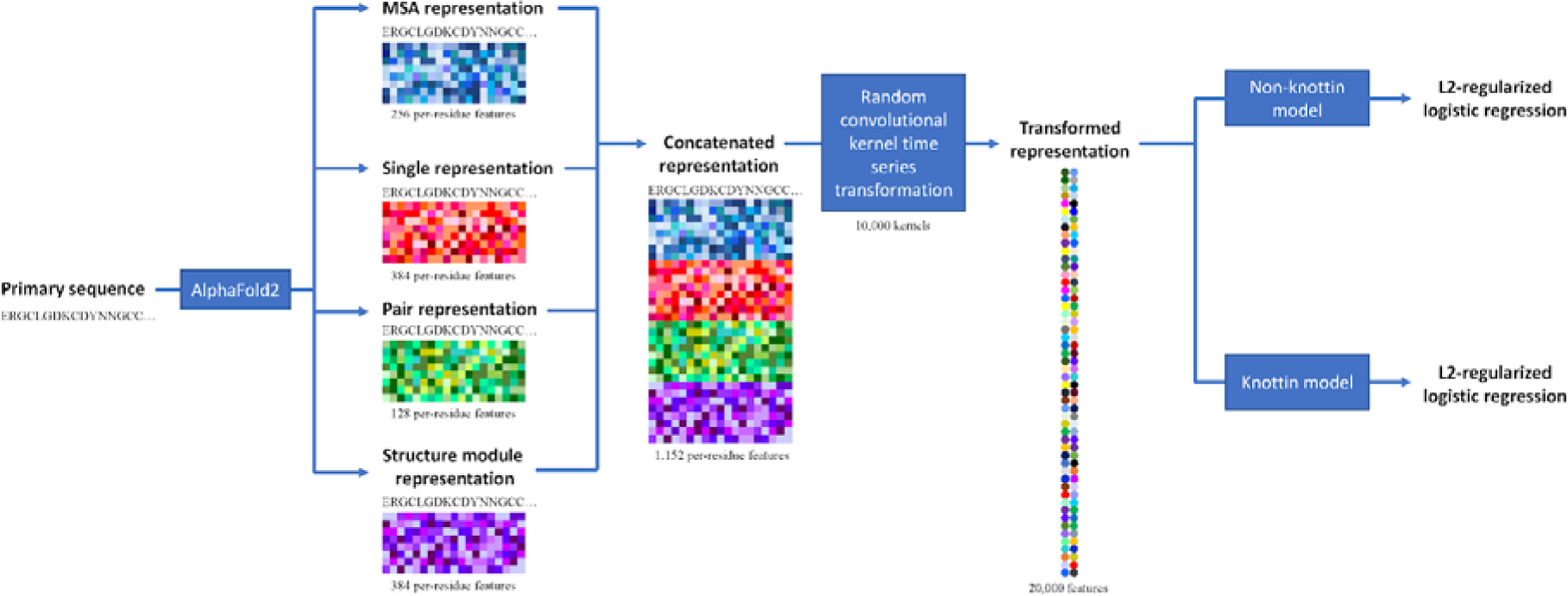
Machine learning architecture diagram of CysPresso. The primary sequence of the CDP is used to generate MSA, single, pair, and structure module AlphaFold2 representations. The four representations are concatenated, and a time series transformation utilizing random convolutional kernels is carried out on the concatenated representation. The transformed representation is then used to predict expressibility using L2-regularized logistic regression machine learning models for knottin and non-knottin CDPs.

**Table 2.** Classifier performance metrics for CysPresso.

| Measure | Non-knottin CDP model | Knottin model |
| --- | --- | --- |
| AUC | 0.798 | 0.852 |
| Sensitivity | 0.853 | 0.764 |
| Specificity | 0.584 | 0.812 |
| Precision | 0.743 | 0.783 |
| Accuracy | 0.742 | 0.789 |
| F1 Score | 0.794 | 0.773 |

The performance of the non-knottin and knottin CysPresso models were evaluated by leave-one-out cross-validation.

### Model interpretability

We then posed the question: Which features are most important for determining expressibility in non-knottin and knottin CDPs? To answer this, we calculated random forest feature contributions by SHAP analysis [31]. Surprisingly, this analysis revealed that the region most important for the prediction of expressibility in non-knottins was near the C-terminus, whereas amino acid residues closer to the N-terminus were deemed more important for predicting expressibility in knottins (Fig. 4A). We then examined the relative importance of the different AlphaFold2 representations in the prediction of expressibility and found that the structure module representation was most important for predicting non-knottin expressibility, while the abstract single representation was most important for predicting knottin expressibility (Fig. 4B). This difference provides a strong rationale for partitioning the dataset into non-knottin and knottin subgroups for improved model performance.

**Fig 4.**
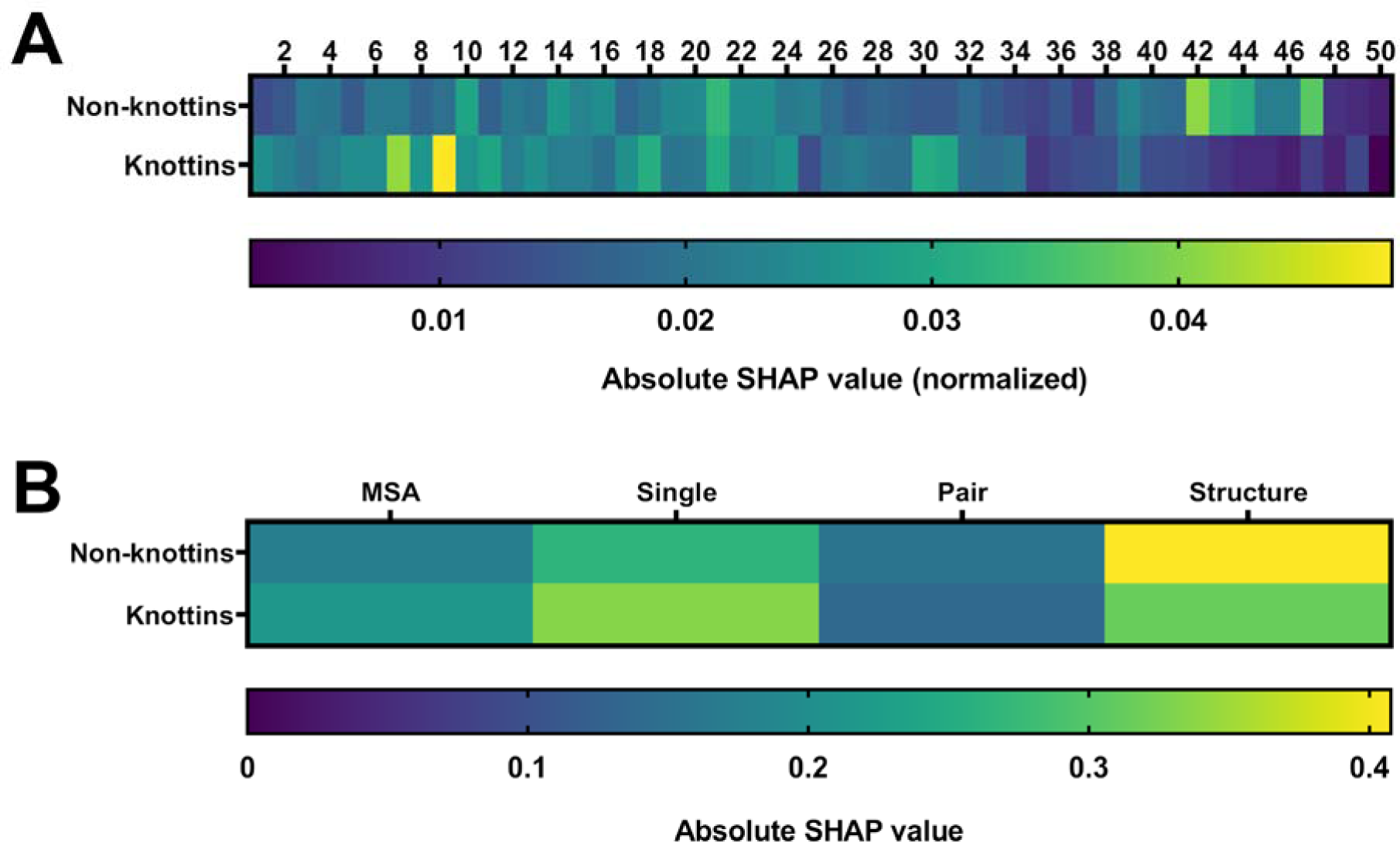
Features important for predicting expressibility in non-knottins are different from the features that are important for predicting expressibility in knottin peptides. This figure shows absolute SHAP values, which illustrate to what degree a feature affects prediction, calculated for different AlphaFold2 features. **(A)** The relative importance of amino acid position for prediction of expressibility in non-knottin and knottin peptides. In non-knottins, a region near the C-terminus (position 42 to 47) was identified as playing an important role in determining expressibility. In knottin peptides, the features at amino acid position 7-9 near the N-terminus were most important for expressibility. **(B)** The relative importance of the different AlphaFold2 representations for expressibility prediction of non-knottin and knottin peptides. For non-knottins, the structure representation module representation was most important for prediction of expressibility. On the other hand, in knottin peptides, the abstract single representation was most important for prediction of expressibility.

AlphaFold2 provides a per-residue confidence metric called pLDDT, which estimates how well the structure prediction should agree with experimentally determined structures. To determine whether the confidence of the AlphaFold2 structure prediction was associated with prediction outcome, the average pLDDT in correctly predicted (true positive and true negative) and incorrectly predicted (false positive and false negative) CDPs were plotted in Figure 5A. The lack of a difference between the average pLDDT values of CDPs that were correctly and incorrectly classified suggests that the confidence of the AlphaFold2 structures do not impact expressibility prediction. This finding is further supported by the fact that the non-knottin CDPs in this study had a higher average pLDDT than knottins (Figure 5B), despite the non-knottin model having a lower performance than that of the knottin model.

**Fig 5.**
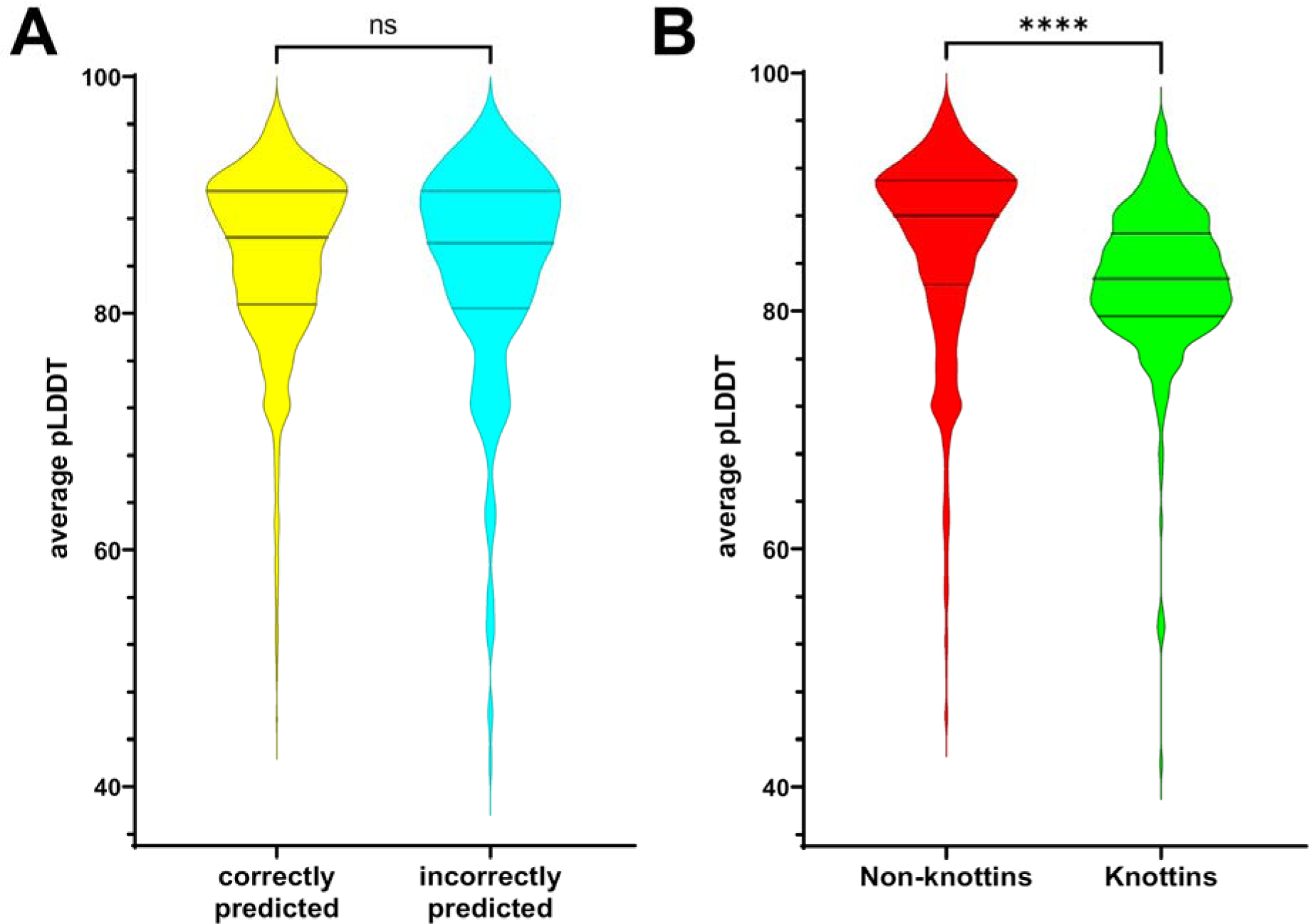
CysPresso outcomes are not biased by pLDDT, a confidence metric of AlphaFold2 predictions. (A) AlphaFold2 average pLDDT values are not significantly different between CDPs that were correctly predicted as expressible and CDPs that were incorrectly predicted. Mann-Whitney test p = 0.512. (B) The average pLDDT values of non-knottin CDPs in the dataset analyzed for this study are significantly higher than those of knottin CDPs. Mann-Whitney test, p < 0.0001. The median and interquartile ranges are indicated by the horizontal lines in the violin plots.

### Comparison to state-of-the-art models

As far as we know, CysPresso is the first machine learning model that predicts the expressibility of recombinant proteins in mammalian cells based on primary sequence. Due to the absence of another model for this classification task, it is challenging to make state-of-the-art comparisons. Recently, a classification model called UniRep-RF was developed using UniRep protein representations obtained from primary sequence to predict recombinant protein expression in *B. subtilis*, where it achieved an AUC of 0.64 [27]. As this model was the closest known comparable model to CysPresso, we obtained UniRep representations [28] for the dataset utilized in this study and evaluated the performance of UniRep-RF and CysPresso using leave-one-out cross-validation (Table 3). Interestingly, UniRep-RF was much better at predicting expressibility of CDPs than it was at predicting recombinant protein expression in *B. subtilis* and achieved an AUC of 0.760. By comparison, CysPresso provided better performance at the task of predicting CDP expressibility from primary sequence and yielded an AUC of 0.816. To test if CysPresso and UniRep-RF were significantly different when evaluated by AUC, we then carried out a non-parametric permutation test and found that CysPresso significantly outperformed UniRep-RF at predicting CDP expressibility (p = 0.0059).

**Table 3.** Comparison of the performance of CysPresso and UniRep-RF at predicting expressibility of CDPs from primary sequence.

| Measure | CysPresso | UniRep-RF |
| --- | --- | --- |
| AUC | 0.816 | 0.760 |
| Sensitivity | 0.818 | 0.786 |
| Specificity | 0.672 | 0.594 |
| Precision | 0.755 | 0.705 |
| Accuracy | 0.753 | 0.700 |
| F1 Score | 0.785 | 0.743 |

The performance of CysPresso and UniRep-RF [27] were evaluated by leave-one-out cross-validation.

## Discussion

### The expressibility of recombinant peptides in human cells can be predicted from primary sequences using machine learning methods

Peptide therapeutics occupy 5% of the global pharmaceutical market, and their approval rate has steadily increased over the last few decades [32]. Presently, peptide therapeutics face two major challenges: the poor yield of peptides from traditional solid-phase synthesis [33] and the intrinsically short plasma half-life of peptides [34]. As a result of recent advances, recombinant peptide production is now a cost effective, simple, and high-yield production platform for peptide drugs [35]. Our tool helps predict which peptides are compatible with recombinant expression, reducing the amount of time-consuming empirical experimentation required to identify expressible CDPs by allowing laboratory efforts to focus on the candidates most likely to express. Furthermore, CysPresso may be useful in the identification of CDPs that are compatible with *in situ* expression technologies such as viral gene therapy and mRNA-based therapeutics, methods that may extend the *in vivo* activity of peptide therapeutics.

To our knowledge there is currently no other model that predicts the expressibility of CDPs (or any difficult-to-express proteins) in mammalian cells based on primary sequence. However, numerous machine learning algorithms have been created for the prediction of heterologous recombinant protein solubility in bacterial expression systems such as *E. coli*. In *E. coli*, overexpression of heterologous proteins can result in protein aggregates and the formation of insoluble inclusion bodies–a phenomenon thought to be a result of misfolding or the formation of intermolecular oligomers. These same factors likely lead to unsuccessful recombinant CDPs expression in mammalian cells. Therefore, prediction of protein solubility in E. coli and predicting CDP expression in mammalian cells is not a dissimilar problem. Recent examples for sequence-based prediction of protein solubility include SoluProt which achieves an AUC of 0.62 [36], SoDoPe which achieves an AUC of 0.71 [37], SKADE which achieves an AUC of 0.82 [38], and DSResSol which achieves an AUC of 0.87 [39]. Our model achieved an AUC of 0.82, which provides comparable performance and it was significantly better at predicting CDP expressibility than UniRep-RF, a *B. subtilis* specific machine learning model for predicting protein expression [27].

### Deep learning protein representations retain information that is complementary to prediction tasks beyond protein structure

In this study, we tested various protein representations and evaluated their usefulness at predicting expressibility of difficult to express peptides. We began our investigation by utilizing neural embeddings from protein language models such as SeqVec and ProteInfer, which are based on statistical models and generated by training large scale sequence datasets. We also utilized AlphaFold2 neural embeddings, which include several representations based on evolutionary, positional, structural, and simulated relationships. AlphaFold2 protein representations provided better predictive performance than the purely language based models. This may be due to the structural information contained within AlphaFold2 representations, as the expressibility of a peptide is determined not only by its amino acid sequence but also by its tertiary and quaternary structure.

While AlphaFold2 accurately predicts protein structures, the suitability of AlphaFold2 neural embeddings for use in general functional prediction tasks is not well established. We find that AlphaFold2 neural embeddings are indeed suitable for general purpose tasks such as the prediction of expressibility of difficult-to-express peptides such as CDPs. Our results agree with work from other groups that have shown that AlphaFold2 produces a general-purpose protein representation that can be used for functional predictions [40]. Though in this study we focused on the expressibility of CDPs, protein representations can theoretically be generated for any class of protein or peptide, thus there is no reason why the methods utilized in this study cannot be generalized to expression datasets of any difficult-to-expression protein. On the same note, protein representations from deep learning protein models, such as those used in this study and others such as RoseTTAFold [32], colabfold [41], openFold [33], proteinBERT [34] and ESMfold [35] have no species bias. Resultantly, there is no reason why the methods in this study cannot be generalized to the prediction of expressibility in any heterologous expression system.

### Different variables determine the expressibility of non-knottin peptides and knottin peptides

Our analysis demonstrated that expressibility is determined by different protein features in knottin and non-knottin CDPs. Knottins are characterized by six cysteine residues making up three interwoven disulfide bridges that form a unique pseudoknot with conserved secondary structure [3]. It is hypothesized that during folding, the C-terminus of the peptide first forms a conserved −hairpin, allowing disulfide bridges to form between cysteines II–V and III–VI. Folding is then completed by the N-terminal loop swinging around to create the cysteine I–IV disulfide bridge [42]. Since our data suggests that the N-terminal region is important for the prediction of expressibility, it is likely that the success of the final N-terminal folding step is required for successful expression of knottin peptides.

Our results also demonstrated that the various AlphaFold2 representations hold differential predictive value. For non-knottins, the structure representation was most important for predicting expressibility, while for knottins, the abstract single representation was most predictive. We speculate that this may be due to the fact that the non-knottin CDPs in this study was a heterogeneous group that included cyclotides, hitchins, growth factor cysteine knots and non-knotted CDPs [17], while the knottin peptides in this study were a homogenous group that all had the conserved inhibitor cysteine knot structure characteristic of knottins. As a result, structurally diverse non-knottin CDPs benefited more from structural information, whereas structural information was less important at providing features that determine expressibility in the structurally related knottins group. In this case, primary sequence relationships captured in the abstract single representation provided more meaningful predictive value.

### Random convolutional kernel time series classification improves protein representation feature performance for prediction of knottin expressibility

The goal of time series classification is to predict the class that a new sample belongs to using labeled ordered training data. While the data used with time series algorithms are typically sequences of discrete-time data, any data registered with some notion of ordering is compatible [43]. As AlphaFold2 and many other protein models such as SeqVec and ProteInfer produce such ordered data (ie. a neural embedding for each amino acid residue), we were able to cast our classification problem as a time series classification problem. We find that applying a time series transformation such as ROCKET improves the capture of ordered features relevant for the prediction of peptide expressibility when examining knottin peptides. Importantly, ROCKET employs random convolutional kernels, which have been shown to be a simple and cost efficient method to capture many features which previously required their own specialized techniques [44]. Thus far, almost all studies that use protein representations for supervised machine learning tasks generate embeddings by taking the average of the pool of activations from the last hidden layer of the model [45]. Our results suggest that the use of time series transformations, in particular–the use of convolutional kernels–provides a better summary of the protein representation by capturing important features that are lost by averaging.

### Study limitations

We analyzed a previously published dataset [17] that examined the expressibility of CDPs in HEK293 cells using a high-throughput Daedalus lentivirus transduction system. In that study, CDPs were considered successfully expressed if the peptide product was free of heterogeneity, as determined by RPC. It is important to note that, while RPC is a rigorous tool for ensuring chemical and conformational homogeneity, it is possible that the expressed products lack biological activity or that heterogeneous products deemed unexpressible may in fact be biologically functional. Additionally, the Daedalus lentiviral expression system that generated the dataset requires proteolytic cleavage to free the peptide from a mouse Ig peptide sequence. Thus, it is possible our results are not generalizable to peptides expressed by other methods.

It is also important to note that the dataset used in this study is likely to be affected by selection biases and coverage bias. For example, our dataset only included CDPs that were between 30 and 50 amino acids long. According to a 2017 analysis [17] of CDPs listed on the RSCB Protein Data Bank [46], non-knottin CDPs ranged from 15 to 81 amino acids in length, and knottin CDPs ranged from 20 to 70 amino acids in length. Resultantly, it is possible that CysPresso may not generalize to CDPs outside of the 30 to 50 amino acid range. Future work should aim to evaluate CysPresso with randomly sampled real-world data.

## Conclusion

In this work, we used supervised machine learning to develop a tool to predict whether a CDP is expressible in a mammalian cell expression system. To accomplish this, we tested the performance of various protein representations and found that AlphaFold2 representations, which are typically used to generate predictions of protein structures, can also be used to predict expressibility. By combining the four AlphaFold2 representations, partitioning the model, and utilizing time series transformation and random convolutional kernels, we further improved the performance of the expressibility classifier to yield an approach with state-of-the-art performance for predicting CDP expressibility in mammalian cells.

## Supplemental Material

Supplemental Table 1: The CDP expressibility dataset utilized in this study.

## Declarations

### Ethics approval and consent to participate

Not applicable. This study does not involve human subjects, vertebrate animals, or human embryonic stem cells.

## Consent for publication

Not applicable. This study does not involve human subjects.

## Availability of data and materials

The datasets generated and/or analyzed during the current study are available in the Hugging Face Repository, https://huggingface.co/datasets/TonyKYLim/CysPresso/tree/main. The code for CysPresso is available as a Colaboratory notebook at https://github.com/Zebreu/cyspresso.

## Author Contributions

**Sébastien Ouellet:** Conceptualization, Methodology, Software, Validation, Formal analysis, Investigation, Data curation, Writing - Original Draft, Writing - Review & Editing, Visualization. **Larissa Ferguson:** Conceptualization, Software, Writing - Review & Editing, Visualization. **Angus Z. Lau:** Investigation, Resources. **Tony K.Y. Lim:** Conceptualization, Methodology, Validation, Formal analysis, Investigation, Writing - Original Draft, Writing - Review & Editing, Visualization.

## Acknowledgements

Not applicable.

## Affiliations

Sébastien Ouellet and Tony K.Y. Lim performed this work as unaffiliated researchers. Tony K.Y. Lim’s current affiliation is the Department of Pharmacology, University of Cambridge.

## References

1. Northfield SE, Wang CK, Schroeder CI, Durek T, Kan M-W, Swedberg JE, et al. Disulfide-rich macrocyclic peptides as templates in drug design. Eur J Med Chem. 2014;77:248–57.

2. Wang CK, Craik DJ. Designing macrocyclic disulfide-rich peptides for biotechnological applications. Nat Chem Biol. 2018;14:417–27.

3. Gracy J, Chiche L. Structure and Modeling of Knottins, a Promising Molecular Scaffold for Drug Discovery. Current Pharmaceutical Design. 2011;17:4337–50.

4. Molesini B, Treggiari D, Dalbeni A, Minuz P, Pandolfini T. Plant cystine-knot peptides: pharmacological perspectives. British Journal of Clinical Pharmacology. 2017;83:63–70.

5. Dongol Y, Cardoso FC, Lewis RJ. Spider Knottin Pharmacology at Voltage-Gated Sodium Channels and Their Potential to Modulate Pain Pathways. Toxins (Basel). 2019;11:E626.

6. Scott A, Weldon S, Taggart CC. SLPI and elafin: multifunctional antiproteases of the WFDC family. Biochem Soc Trans. 2011;39:1437–40.

7. Muratspahi E, Koehbach J, Gruber CW, Craik DJ. Harnessing cyclotides to design and develop novelćpeptide GPCR ligands. RSC Chem Biol. 2020;1:177–91.

8. Gracy J, Le-Nguyen D, Gelly J-C, Kaas Q, Heitz A, Chiche L. KNOTTIN: the knottin or inhibitor cystine knot scaffold in 2007. Nucleic Acids Res. 2008. https://doi.org/10.1093/nar/gkm939.

9. Postic G, Gracy J, Périn C, Chiche L, Gelly J-C. KNOTTIN: the database of inhibitor cystine knot scaffold after 10 years, toward a systematic structure modeling. Nucleic Acids Res. 2018;46 Database issue:D454–8.

10. Kintzing JR, Cochran JR. Engineered knottin peptides as diagnostics, therapeutics, and drug delivery vehicles. Current Opinion in Chemical Biology. 2016;34:143–50.

11. Russo A, Aiello C, Grieco P, Marasco D. Targeting “Undruggable” Proteins: Design of Synthetic Cyclopeptides. Current Medicinal Chemistry. 2016;23:748–62.

12. Visintin M, Melchionna T, Cannistraci I, Cattaneo A. In vivo selection of intrabodies specifically targeting protein–protein interactions: A general platform for an “undruggable” class of disease targets. Journal of Biotechnology. 2008;135:1–15.

13. de Araujo CB, Heimann AS, Remer RA, Russo LC, Colquhoun A, Forti FL, et al. Intracellular Peptides in Cell Biology and Pharmacology. Biomolecules. 2019;9:150.

14. Reinwarth M, Glotzbach B, Tomaszowski M, Fabritz S, Avrutina O, Kolmar H. Oxidative Folding of Peptides with Cystine-Knot Architectures: Kinetic Studies and Optimization of Folding Conditions. ChemBioChem. 2013;14:137–46.

15. Reinwarth M, Nasu D, Kolmar H, Avrutina O. Chemical Synthesis, Backbone Cyclization and Oxidative Folding of Cystine-knot Peptides — Promising Scaffolds for Applications in Drug Design. Molecules. 2012;17:12533–52.

16. Rivera-de-Torre E, Rimbault C, Jenkins TP, Sørensen CV, Damsbo A, Saez NJ, et al. Strategies for Heterologous Expression, Synthesis, and Purification of Animal Venom Toxins. Front Bioeng Biotechnol. 2022;9:811905.

17. Correnti CE, Gewe MM, Mehlin C, Bandaranayake AD, Johnsen WA, Rupert PB, et al. Screening, large-scale production and structure-based classification of cystine-dense peptides. Nat Struct Mol Biol. 2018;25:270–8.

18. Crook ZR, Sevilla GP, Friend D, Brusniak M-Y, Bandaranayake AD, Clarke M, et al. Mammalian display screening of diverse cystine-dense peptides for difficult to drug targets. Nat Commun. 2017;8:2244.

19. Heinzinger M, Elnaggar A, Wang Y, Dallago C, Nechaev D, Matthes F, et al. Modeling aspects of the language of life through transfer-learning protein sequences. BMC Bioinformatics. 2019;20:723.

20. Sanderson T, Bileschi ML, Belanger D, Colwell LJ. ProteInfer: deep networks for protein functional inference. 2021;:2021.09.20.461077.

21. Jumper J, Evans R, Pritzel A, Green T, Figurnov M, Ronneberger O, et al. Highly accurate protein structure prediction with AlphaFold. Nature. 2021;596:583–9.

22. Dempster A, Petitjean F, Webb GI. ROCKET: exceptionally fast and accurate time series classification using random convolutional kernels. Data Min Knowl Disc. 2020;34:1454–95.

23. The UniProt Consortium. UniProt: the universal protein knowledgebase in 2021. Nucleic Acids Research. 2021;49:D480–9.

24. Sanderson T, Bileschi ML, Belanger D, Colwell LJ. ProteInfer, deep neural networks for protein functional inference. eLife. 2023;12:e80942.

25. Breiman L. Random Forests. Machine Learning. 2001;45:5–32.

26. Pedregosa F, Varoquaux G, Gramfort A, Michel V, Thirion B, Grisel O, et al. Scikit-learn: Machine Learning in Python. Journal of Machine Learning Research. 2011;12:2825–30.

27. Martiny H-M, Armenteros JJA, Johansen AR, Salomon J, Nielsen H. Deep protein representations enable recombinant protein expression prediction. Computational Biology and Chemistry. 2021;95:107596.

28. Alley EC, Khimulya G, Biswas S, AlQuraishi M, Church GM. Unified rational protein engineering with sequence-based deep representation learning. Nat Methods. 2019;16:1315– 22.

29. Goodfellow I, Bengio Y, Courville A. Deep Learning. MIT Press; 2016.

30. Demšar J. Statistical Comparisons of Classifiers over Multiple Data Sets. J Mach Learn Res. 2006;7:1–30.

31. Lundberg SM, Lee S-I. A Unified Approach to Interpreting Model Predictions. In: Advances in Neural Information Processing Systems. Curran Associates, Inc.; 2017.

32. Muttenthaler M, King GF, Adams DJ, Alewood PF. Trends in peptide drug discovery. Nat Rev Drug Discov. 2021;20:309–25.

33. Isidro-Llobet A, Kenworthy MN, Mukherjee S, Kopach ME, Wegner K, Gallou F, et al. Sustainability Challenges in Peptide Synthesis and Purification: From R&D to Production. J Org Chem. 2019;84:4615–28.

34. Kontermann RE. Strategies for extended serum half-life of protein therapeutics. Curr Opin Biotechnol. 2011;22:868–76.

35. Ajingi YS, Rukying N, Aroonsri A, Jongruja N. Recombinant Active Peptides and their Therapeutic Functions. Current Pharmaceutical Biotechnology. 2022;23:645–63.

36. Hon J, Marusiak M, Martinek T, Kunka A, Zendulka J, Bednar D, et al. SoluProt: prediction of soluble protein expression in Escherichia coli. Bioinformatics. 2021;37:23–8.

37. Bhandari BK, Gardner PP, Lim CS. Solubility-Weighted Index: fast and accurate prediction of protein solubility. Bioinformatics. 2020;36:4691–8.

38. Raimondi D, Orlando G, Fariselli P, Moreau Y. Insight into the protein solubility driving forces with neural attention. PLoS Comput Biol. 2020;16:e1007722.

39. Madani M, Lin K, Tarakanova A. DSResSol: A Sequence-Based Solubility Predictor Created with Dilated Squeeze Excitation Residual Networks. IJMS. 2021;22:13555.

40. Hu M, Yuan F, Yang KK, Ju F, Su J, Wang H, et al. Exploring evolution-based & −free protein language models as protein function predictors. 2022.

41. Mirdita M, Schütze K, Moriwaki Y, Heo L, Ovchinnikov S, Steinegger M. ColabFold: making protein folding accessible to all. Nat Methods. 2022;19:679–82.

42. McCarthy S, Robinson J, Thalassinos K, Tabor AB. A Chemical Biology Approach to Probing the Folding Pathways of the Inhibitory Cystine Knot (ICK) Peptide ProTx-II. Frontiers in Chemistry. 2020;8:228.

43. Gamboa JCB. Deep Learning for Time-Series Analysis. 2017.

44. Wang Z, Yan W, Oates T. Time Series Classification from Scratch with Deep Neural Networks: A Strong Baseline. 2016.

45. Yang KK, Wu Z, Bedbrook CN, Arnold FH. Learned protein embeddings for machine learning. Bioinformatics. 2018;34:2642–8.

46. Burley SK, Bhikadiya C, Bi C, Bittrich S, Chen L, Crichlow GV, et al. RCSB Protein Data Bank: powerful new tools for exploring 3D structures of biological macromolecules for basic and applied research and education in fundamental biology, biomedicine, biotechnology, bioengineering and energy sciences. Nucleic Acids Research. 2021;49:D437–51.

